# Does Antibody Stabilize the Ligand Binding in GP120 of HIV-1 Envelope Protein? Evidence from MD Simulation

**DOI:** 10.1101/595413

**Authors:** Vishnudatt Pandey, Rakesh Kumar Tiwari, Rajendra Prasad Ojha, Kshatresh Dutta Dubey

## Abstract

CD4-mimetic HIV-1 entry inhibitors are small sized molecules which imitate similar conformational flexibility in gp120 as CD4 receptor, the mechanism of the conformational flexibility instigated by these small sized inhibitors, however, is little known. Likewise, the effect of the antibody on the function of these inhibitors is also less studied. In this study, we present a thorough inspection of the mechanism of the conformational flexibility induced by a CD4-mimetic inhibitor, NBD-557, using Molecular Dynamics Simulations and free energy calculations. Our result shows a functional importance of Asn239 in substrate instigated conformational dynamics in gp120. The MD simulations of Asn239Gly mutant provide a less dynamic gp120 in the presence of NBD-557 without incapacitating the binding enthalpy of NBD-557. The MD simulations of complex with the antibody clearly shows the enhanced affinity of NBD-557 due to the presence of the antibody which is in good agreement with experimental Isothermal Titration Calorimetry results (*Biochemistry* **2006,** *45*, 10973-10980).

## 1. Introductions

The pathogenic importance of human immunodeficiency virus type-1 (HIV-1) is well known.^1-3^ The HIV-1 infection is caused by the fusion of viral and host membranes in many sequential steps which are initiated by the non-covalent attachment of two regulatory subunits of HIV-1 envelope protein, gp120 and gp41.^4^ A schematic view of these sequential steps can be comprehended by scheme1.

**Scheme1.**
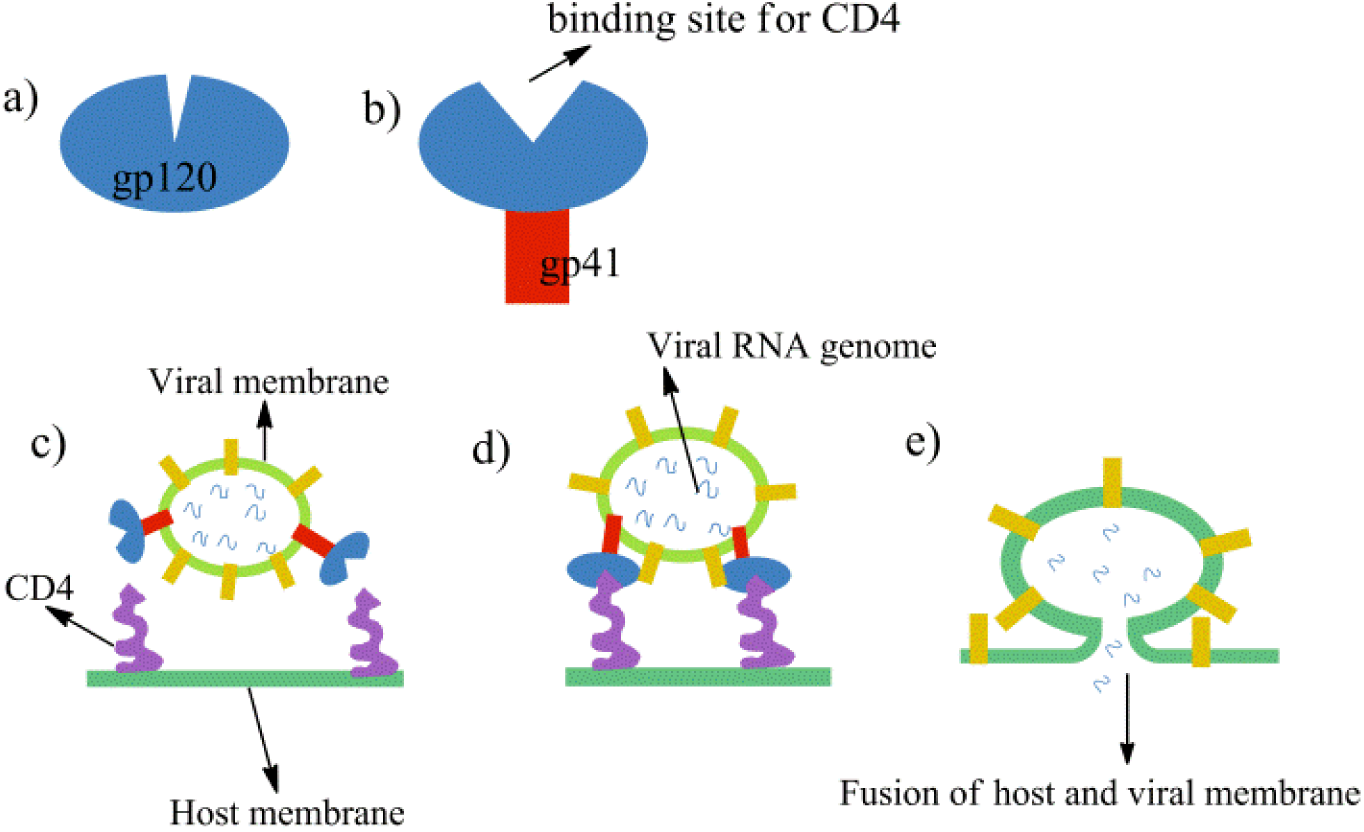
A schematic representation of HIV-1 entry and fusion of virion into host membrane.

Initially, each unit of gp120 protein are arranged into homotrimeric fashion and form a closed cavity mushroom head-type system as shown in **a**, while the non-covalent binding of the gp41 protein with gp120 triggers some conformational changes that open a binding cavity for CD4 receptors and forms a complete mushroom shape, shown in **b**.^5-6^ The HIV-1 virion contains the RNA genome encapsulated by viral membrane where gp120.gp41 complexes are attached to the membrane surface and are exposed to the CD4 protein receptors of host membrane as shown in **c**.^7-8^ The docking of the virion via gp120 to the host membrane gives rise to the conformational rearrangement in the envelope protein that pulls viral membrane close to the CD4 subunits of host membrane,^9^ shown in **d**. The binding of CD4 receptors via gp120 protein leads to more conformational changes and finally the fusion of host and viral membrane takes place, shown in **e**. As clear from the scheme 1, the gp120 is central to viral entry via binding to gp41 and CD4; it turns out to be the hot topic to target HIV-1 infections. The structural topography of gp120 revealed by X-ray crystallography shows two domains mixed of α- and β-motifs constitute the structure of gp120: the inner domain (residues 1-28, 44-95, 289-300) which interacts with gp41 protein, and outer and less conserved domain which interacts with CD4 and co-receptor (CR) proteins (see Figure 1 for the structure of gp120)^10-11^. Since the binding of gp120 with CD4 receptor proteins is the crucial step during HIV-1 infections, therefore, the site of CD4 binding in gp120 (scheme 1b) becomes a productive target for HIV-1 inhibition.^12-15^The details and the recent development of these HIV-1 entry inhibitors targeting the gp120 proteins can be found in some nice review articles where we observe majority of the HIV-1 entry inhibitors are either BMS-378806 and its analogous or NBD-556 and its derivative.^16-18^ Interestingly, these two inhibitors target same receptor but their mechanism is entirely different. An experimental study of Schön et al shows that the binding mode and antiviral properties of these inhibitors are governed by their binding thermodynamics.^19^ In same study authors show that BMS-378806 has very weak binding enthalpy and does not compete with CD4-coreceptor, on the other hand NBD is CD4-coreceptor competitive inhibitor and reflects similar conformational flexibility and binding thermodynamics as CD4-coreceptor. However, *the rationale that how binding of small molecule such as NBD instigates the conformational flexibility in gp120 remains an enigma. The current study provides a theoretical explanation of the mechanism of the conformational dynamics instigated by inhibitor.*

**Figure 1.**
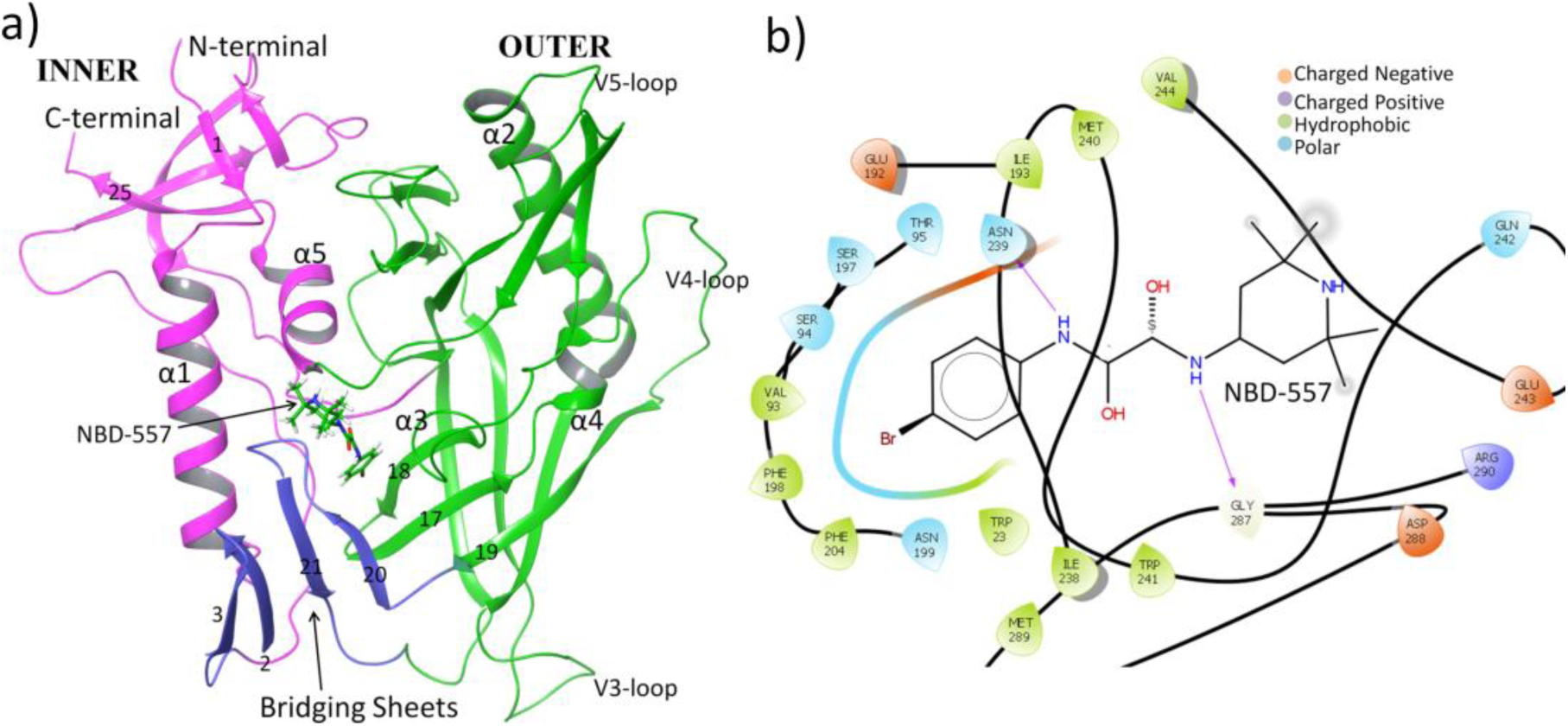
a) 3D structure of gp120 with structural topology. Inner domain is shown by pink, outer domain is shown by green while bridging sheets are shown by blue colors. Numbering of β-sheets is taken from ref.11 and only few of them are shown to maintain minimalism in the figure, b) 2D ligand interaction diagram of gp120 from the crystal structure^20^, H-bonds are shown by pink arrows.

The instability of gp120 protein, when separated from its membrane anchored part, is a fundamental problem in getting the crystal structure of gp120.^4^ Therefore most of the structures of gp120 were obtained with core domain bound antibody even in the presence of HIV-1 entry inhibitors. *This often raises a possibility that the binding of HIV-1 inhibitor in presence of antibody may influence the interactions of inhibitor. Does antibody presence affect the mechanism of HIV-1 inhibitor? If, yes then how? This study provides a valuable insight on these questions using Molecular Dynamics (MD) Simulations and free energy calculations.* The study of Schön et al provides a little hint that the presence of antibody may enhance the ligand binding and thus efficacy of the inhibitor, however, a comprehensive investigation is still missing. We have provide a mechanistic insight and a comprehensive study for the affect of antibody on the binding of HIV-1 entry inhibitors using sufficiently long molecular dynamics (MD) simulations with a typical example of NBD-557 with gp120. In order to achieve such study, we firstly emphasize the NBD-557 interactions and its binding thermodynamics without antibody and then compared the NBD-557 interaction and it’s binding in the presence of antibody.

## 2. Methods

### 2.1 System Setup

The initial structure was taken from protein data bank (PDB id: 4DVR)^20^ which contains gp120 core, ligand NBD-557, and antibody Fab 48d. Since we have focused on the role of antibody on NBD-557 binding, we prepared two systems; one by extracting antibody and constructing a complex of ligand NBD-557 with gp120 only, and second as ligand NBD-557 with gp120 and antibody. The protonation state of Histidine residues was calculated by propka 3.0.^21^ The pKa value for His54 was 6.72, therefore we kept it at doubly protonated state i.e. HIP, and rests of the Histidines were kept at either HIE (epsilon protonation) or HID (delta protonation) state depending upon the H-bond possibility with nearby residues. The missing hydrogens were prepared with Leap module of Amber 14 with Amber ff14SB forcefield parameters.^22^ Since the parameters for NBD-557 were not available in Amber library, we used antechamber module of Amber to generate the parameters for NBD-557. The partial atomic charges and missing parameters for the NBD-557 were obtained from the RESP^23-24^ (Restrained electrostatic potential) method using HF/6-31G* geometry optimization by Gaussian 09 package. We used five and two Cl^-^ ions to neutralize the NBD-557 with gp120 only complex and the NBD-557 with gp120 and antibody complex, respectively. Finally, the resulting systems were solvated in a rectangular box of TIP3P^25^ waters extending up to minimum cutoff of 10 Å from the protein boundary.

### 2.2. MD Simulations

After proper parameterizations and setup, the resulting system’s geometries were minimized (5000 steps for steepest descent and 10000 steps for conjugate gradient) to remove the poor contacts and relax the system. The systems were then gently annealed from 10 to 300 K under canonical ensemble for 50 ps with a weak restraint of 5 kcal/mol/Å^2^. Subsequently, the systems were maintained for 1 ns of density equilibration under isothermal-isobaric ensemble at target temperature of 300K and the target pressure of 1.0 atm using Langevin-thermostat^26^ and Berendsen barostat^27^ with collision frequency of 2 ps and pressure relaxation time of 1 ps, with a weak restraint of 1 kcal/mol/Å^2^. Thereafter, we removed all restraints applied during heating and density dynamics and further equilibrated the systems for ∼3 ns to get well settled pressure and temperature for conformational and thermodynamical analyses. This was followed by a productive MD run of 100 ns for each system. During all MD simulations, the covalent bonds containing hydrogen were constrained using SHAKE,^28^ and particle mesh Ewald (PME)^29^ was used to treat long-range electrostatic interactions. We used an integration step of 2 fs during the entire simulations. All MD simulations were performed with GPU version of Amber 14 package.^30^

### 2.3 Clustering of the Trajectories

The energetic of ligand binding using end point method such as MMGB/SA (Molecular Mechanical Generalized Boltzmann/Surface Area) depends upon the selection of MD frames.^31^ In this reference, clustering of trajectories provides an accurate way to represent the most statistically significant structures during MD sampling. Therefore, to facilitate most persistent conformations of the system during simulation, we performed clustering of MD trajectories using hieragglo algorithm implemented in Cpptraj of the Amber MD package for all complexes. The clustering was done for the protein ligand complexes. In each case 10 clusters were generated. The results of the clustering are shown in Supplementary material.

### 2.4 Free energy Calculations

As stated by Schön et al, the antiviral properties of HIV-1 entry inhibitors are reflected by the thermodynamics of these inhibitors, we used MM-GB/SA method^32-33^ for the calculations of thermodynamic parameters and free energy of binding. The principles of these methods are well established^34-35^ and have been successfully applied for the class II fusion viruses in the previous studies^36-38^. Note that all energetics are calculated for the most populated frames of MD trajectories obtained from cluster analysis as described above. All water molecules and the chloride ions were stripped from the trajectory prior to the MMPBSA analysis. The dielectric constant for the solute and the surrounding solvents were kept 1 and 80 respectively. The MMPBSA calculations were performed on the most populated trajectories obtained from cluster analysis. All analysis of trajectories were done with the Cpptraj module of Amber14. VMD 1.6.7^39^, Chimera-1.5^40^ graphical programs were used for the visualization. The hydrogen bonds and its occupancies were calculated by VMD for production trajectories where we kept donor-acceptor distance as 3.0 Å and angle cut-off as 20°.

## 3. Results and Discussion

We start with the conformational properties of gp120 complex with ligand NBD-557 when antibody is not binding; thereafter we discuss the conformational properties of gp120 complex with ligand NBD-557 when antibody is binding.

### 3.1 MD simulation of gp120 with NBD-557

The RMS deviations (root mean square) with simulation time for the MD simulation of gp120 with NBD-557 are constant which shows a converged MD trajectory (see Figure S1 in supplementary materials). We also simulated gp120 only in absence of NBD-557 to study the effect of ligand binding. The RMS deviations of gp120 in presence of ligand (NBD-557) is slightly more converged *vis-a-vis* gp120 in absence of ligand, which shows ligand binding stabilizes the simulation.

The root mean square fluctuations (RMSF) of gp120 with and without NBD-557, as shown in Figure 2, very clearly emphasize the effect of ligand binding. We can see that the conformational flexibility of gp120 is very high in presence of ligand *vis-a-vis* when ligand is not binding. More importantly, the functional elements (the inner domain that interacts with gp41, the outer domain that interacts with CD4 and the CR proteins of the target cells) show highest flexibility. The increased conformational flexibility at gp41 and CD4+CR binding interfaces due to inhibitor binding substantiates the fact that NBD-557 is a CD4-mimetic inhibitors as similar flexibility in gp120 is supposed to be induced by binding of CD4 co-receptor.^5^ The increased flexibility due to NBD-557 may raise an obvious question that how the binding of a small molecule binding can trigger the conformational dynamics in gp120 and what is the driving force for this phenomenon? Therefore, we thoroughly monitored the interactions of NBD-557 with active site residues and *found that it is interactions between NBD-557 with Asn 239 which drives the flexibility of gp120 with course of simulations* (see Figure 3).

**Figure 2.**
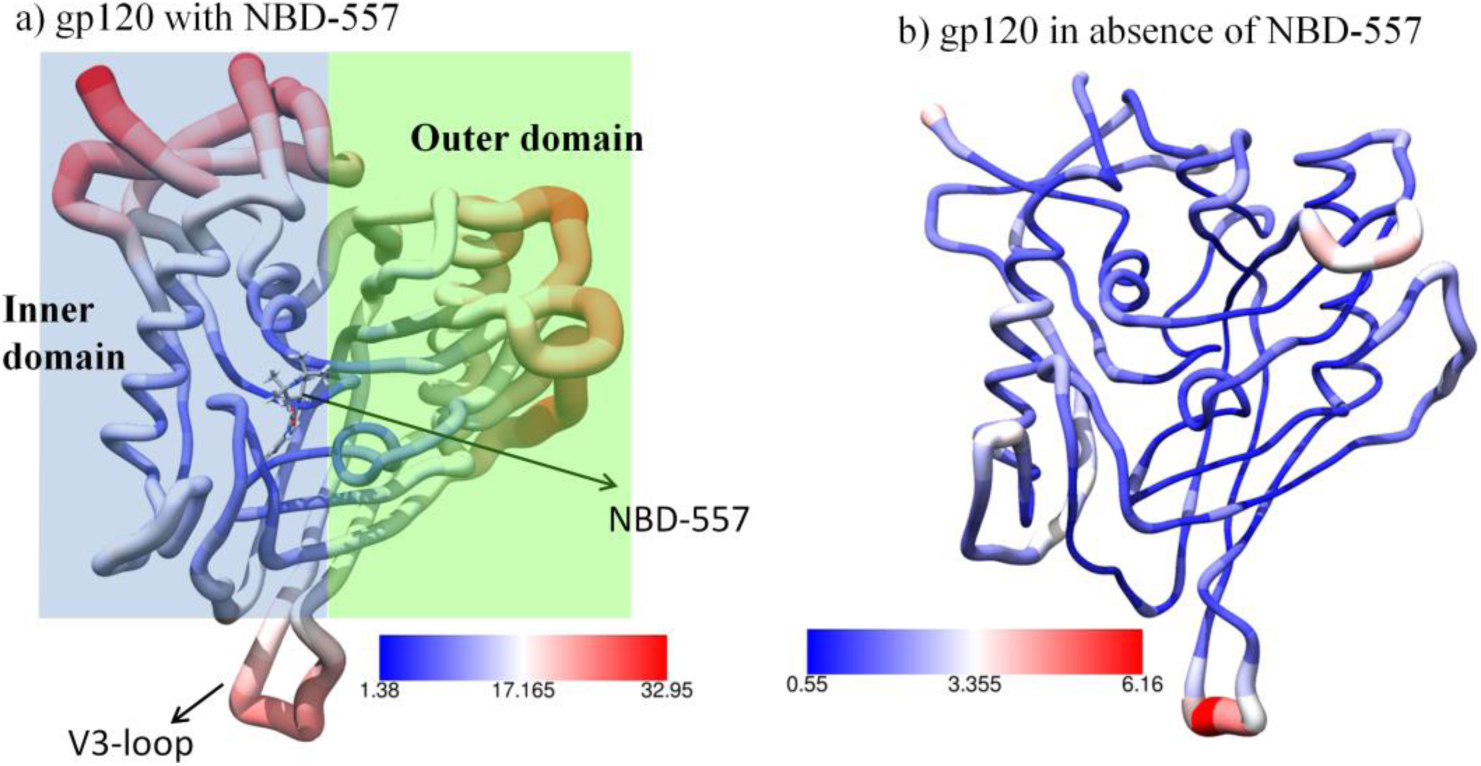
The root mean square fluctuations for gp120 shown in worm model. The thickness of worm radius represents the value of fluctuations, a) with NBD-557, the inner and outer domains are highlighted by blue and green transparent rectangles b) without drug NBD-557. The values of RMSF (Å^2^) are shown by colour keys in each figure.

**Figure 3.**
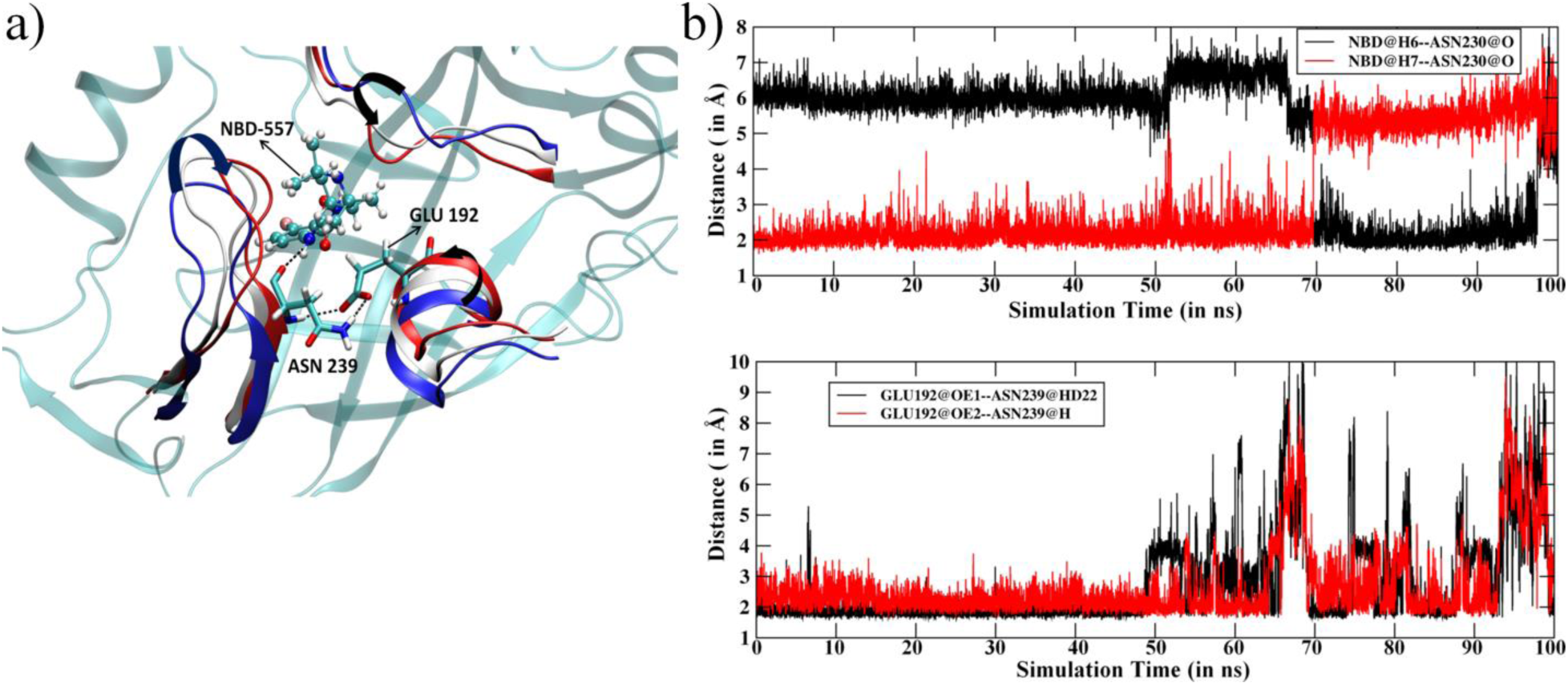
a) Key residues interacting with NBD-557. Notice the movement of structural motifs surrounding ligand indicated by arrows (blue represent initial, white shows mid and red shows the orientations at last step of MD simulations b) the distance *vs* simulation time for 100 ns of MD simulation for key H-bonds (top) and the distance between Glu192 and Asn239 (bottom). Note a constant and strong interaction of NBD-557 with Asn239.

The ligand NBD-557 contains two amine hydrogens situated at two different positions (see Figure 1b) which are prone to make hydrogen bond with backbone oxygen of Asn239. This Adn239 forms a strong hydrogen bond with another residue Glu192. Interestingly, these two residues are part of the inner and outer domains respectively; therefore, NBD-557 acts as a bridge between two functional subunits of gp120. During the course of MD simulations, the amine hydrogens of NBD-557 exchanges their positions to form a strong bond with backbone oxygen of Asn239 (note the exchange of H-bonding from 70-100 ns of MD simulations and a strong correlation between the upper and lower plots during 50-100 ns of simulations; we can easily see that the fluctuation in lower graph is strongly correlated with H-bond exchange in upper plot) which in turns produces a local dynamics in Asn239 and thus in Glu192. Interestingly, Asn239 and Glu192 are residues situated at bridging interfaces between inner and outer domains and a disturbance at this site produces fluctuations at inner and outer domain end as seen in Figure 2a. For further validation of this mechanism, we performed another MD simulation of gp120 in absence of ligand and found a drastic decrement in thermal fluctuations in gp120 (see Figure 2b). In this case, the binding between the Asn239 and Glu192 is rather weak due to absence of ligand which might be reason for the conformational elasticity of gp120 (see Figure S2 for Asn239-Glu192 distance). The lack of thermal fluctuations in absence of NBD-557 strongly supports the *ligand instigated dynamics in gp120 which is factually caused by the interaction between NBD-557 and Asn239.*

### 3.2. Thermo-chemistry of gp120 complex with NBD-557

Schön et al reported the thermodynamics of binding of NBD-556 and NBD-557 with gp120 using ITC (Isothermal Titration Calorimetry).^19^ They found the change in enthalpy during binding of NBD-556 (same as NBD-557) as −24.50 kcal/mol, and can be used to benchmark our simulated structure. Therefore we calculated the change in enthalpy using the MMGBSA method for NBD-557 binding as shown in Table 1. The computed change of enthalpy is −25.16±3.24 kcal/mol and is in good agreement with experimental results.^19^ The theoretical calculation shows that the van der Waals interactions (−36.94 kcal/mol) plays a dominating role in enthalpic change during binding. The electrostatic contribution is rather weak and is quite expected as NBD-557 forms just two hydrogen bonds with Glu 192 and Asn239. Furthermore, the interaction energy due to solvation has unfavourable contribution.

**Table 1.**
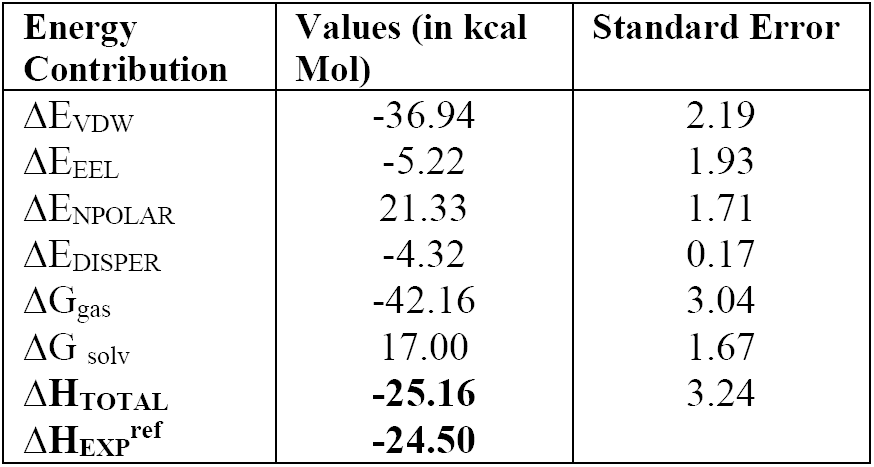
Energy contribution when GP120 bind with NBD in absence of antibody. Here, ΔE_VDW_ =change in van der Waals interaction on ligand associations, ΔE_EEL_ =change in electrostatic interaction on ligand associations, ΔE_polar_ = change in polar interaction on ligand associations, ΔE_nonpolar_ = change in nonpolar interaction on ligand associations. ΔG_gas_ = ΔE_VDW_ + ΔE_EEL_, ΔG_solv_ =ΔE_polar_ +ΔE_nonpolar_, ΔG_binding_=ΔG_gas_ +ΔG_solv_

| Energy Contribution | Values (in kcal Mol) | Standard Error |
| --- | --- | --- |
| $\Delta E_{\text{VDW}}$ | -36.94 | 2.19 |
| $\Delta E_{\text{EEL}}$ | -5.22 | 1.93 |
| $\Delta E_{\text{NPOLAR}}$ | 21.33 | 1.71 |
| $\Delta E_{\text{DISPER}}$ | -4.32 | 0.17 |
| $\Delta G_{\text{gas}}$ | -42.16 | 3.04 |
| $\Delta G_{\text{solv}}$ | 17.00 | 1.67 |
| $\Delta H_{\text{TOTAL}}$ | **-25.16** | 3.24 |
| $\Delta H_{\text{EXP}}^{\text{ref}}$ | **-24.50** | |

To further emphasize the gp120-ligand interactions, we calculated the residue interaction map as shown in Figure 4. The residue interaction map calculated by MMGBSA is very informative as non-polar and hydrophobic interactions cannot be analyzed similar to H-bond and salt-bridges. We can see that Glu192 and Asn239 interact strongly with NBD-557 due to H bond as shown in Figure 4. The ligand is mostly in contact with several hydrophobic residues such as Trp23, Val93, Ile193 and 238, Phe198 and 204, Met240 and 289, and Gly286 and 287. These hydrophobic and non-polar residues contribute significantly in total favourable change of enthalpy during binding. Interestingly similar interactions were predicted in crystal structure of NBD-557 with gp120,^20^ therefore the calculated residue interactions map provides a theoretical validation and shows the stability of such interactions during 100ns of MD simulations.

**Figure 4.**
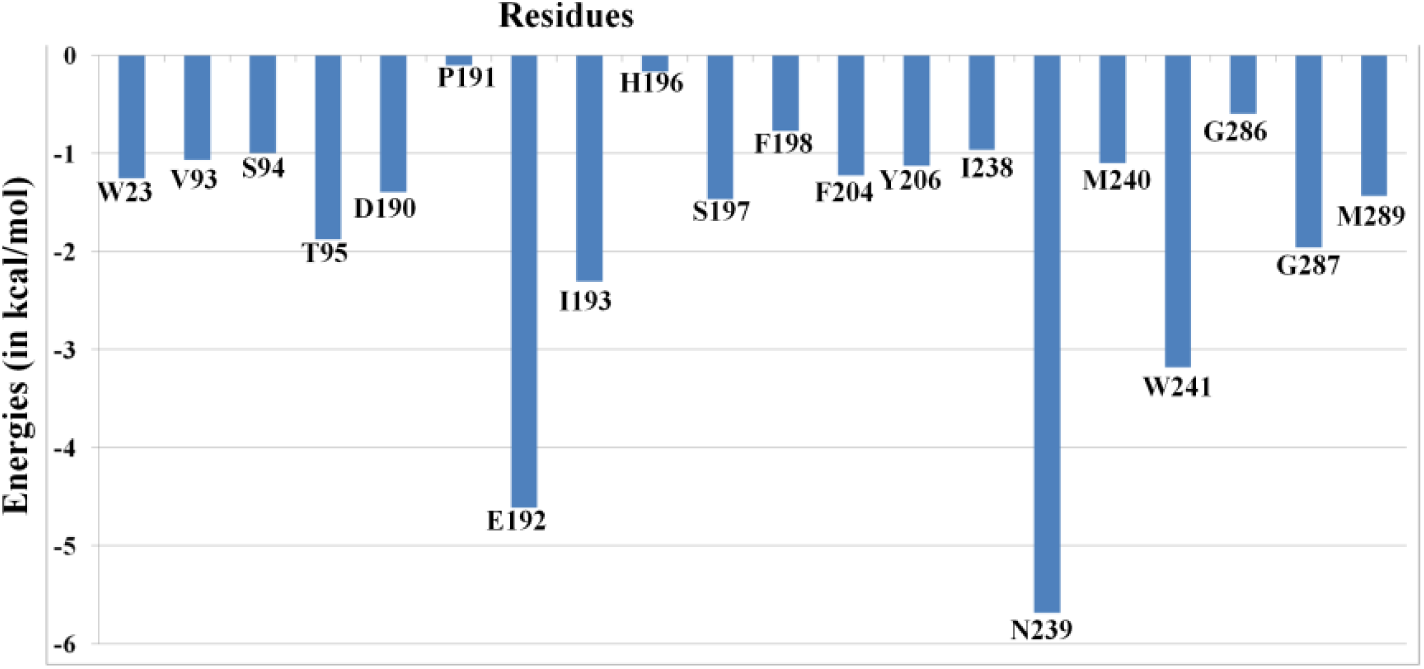
The residue interaction map for NBD with gp120.

### 3.3. Dynamics of the Asn239Gly Mutant

Since NBD-557 interacts strongly with Asn239, we mutated Asn239 to Gly239 to monitor the effect of mutation on the gp120 complex. Interestingly, the ligand remains quite stable. In the Asn239Gly mutant, the one of amine hydrogen (H1--N) of NBD-557 forms a very strong hydrogen bond with backbone oxygen of Glycine and doesn’t show the switching of position as seen in WT complex (see Figure 5a,b). However, the Gly239 interaction with Glu192 is not as strong as in WT complex. The change in enthalpy of binding is more favourable *vis-a-vis* WT complex which clearly indicates that NBD-557 is a strong binder for the mutant complex (see Table S1).

**Figure 5.**
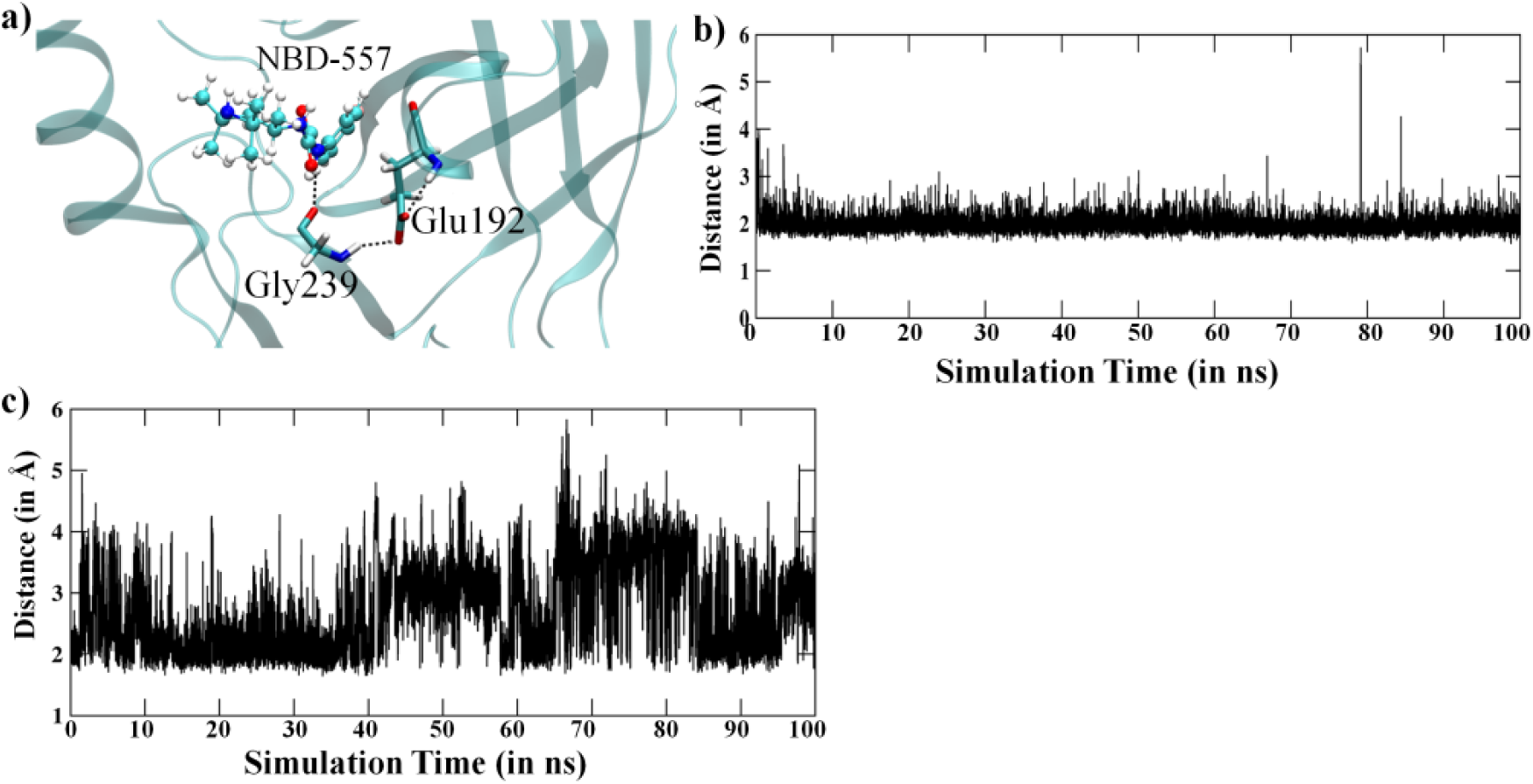
Results from the MD of N239G mutant. a) NBD-557 binding in the gp120 binding pocket, b) The distance *vs* simulation time for H-bond between amine hydrogen of NBD-557 with backbone oxygen of Gly239, c) distance vs time for H-bond between Gly239 and Glu192.

### 3.4. Effect of antibody binding on gp120 interaction with NBD-557

The study by Schön et al indicates that the binding of antibody increases the binding of NBD-556 and NBD-557. They found that the enthalpy change on binding of ligand increases from −24.5 kcal/mol to −28.9 kcal/mol in presence of antibody. Therefore we performed a 200 ns long MD simulation of gp120 complex with NBD in the presence of antibody (data are reported for last 100 ns for a consistency with N239G and gp120 with NBD-557 only simulations) and calculated the change in enthalpy on NBD-557 binding. Our theoretical calculations strongly support the finding of Schön et al that antibody presence enhances the binding affinity of NBD-557. A comparison of RMS deviations for NBD-557 in WT complex, R339G mutant and gp120 complex with NBD is shown in Figure 6. We can easily notice the stability of the NBD-557 in gp120 with antibody relative to WT and mutant. The change in enthalpy of binding is shown in Table 2 which quantitatively supports the enhanced binding affinity of drug due to antibody bonding. To decipher the rationale for the increased affinity due to antibody, we thoroughly studied the interactions between gp120 and antibody. We notice that residues 334-367 of antibody are proximal to gp120 interface and may interact with gp120. The interface between antibody and gp120 are rich of acidic residues such as Asp310, Asp334, Asp331, Asp334 at antibody site, and Arg149, Arg233, Lys235 and Lys246 at gp120 site which provide stability to the gp120 and antibody complex. The interaction energy of these residues, calculated by residue decomposition of MMPBSA is shown in Table 3. These data quantitatively confirms that there are very strong acid-base pair interactions between gp120 and antibody interface such as Asp331-Arg233, Asp334-Arg233, Asp334-Lys235, Asp336-Lys235 and Asp336-Lys246 (See Figure 7 and Table 3).

The results shown above provide compelling evidence that the presence of antibody in gp120 stabilizes the complex via acid-base pair interactions at binding interface, however, whether the antibody affects the drug binding is still unknown. Therefore, we calculated the change in enthalpy upon ligand binding in presence and absence of antibody (See Table 1 for NBD-557 binding enthalpy in absence of antibody). Our calculations, shown in Table 4, show that the presence of antibody not only stabilizes the gp120 complex via acid base pair interactions, but also increases the binding of ligand. We can see that in absence of antibody, the change in enthalpy due to ligand binding is −25.16 kcal/mol while in presence of antibody it significantly increases to −33.57 kcal/mol. To investigate the rationale for the increased affinity we thoroughly monitored the MD trajectories and the residues of gp120 participating in ligand binding and antibody-gp120 binding. Figure 8a shows a comparative interaction map due to antibody binding. It is quite clear that the binding of antibody increases the interactions with NBD for some residues *e.g.* Met240, Trp241, Val244, Gly287, Asp288 which provide stability to the binding of NBD-557. As Figure 8b shows these residues reside on the bridging sheet 20 (see Figure 1b and ref. 11 for structural topology) which is interfacial loop interacting to antibody, therefore these residues shows conformational rearrangement which might cause increased interactions with NBD on antibody binding.

**Table 2.**
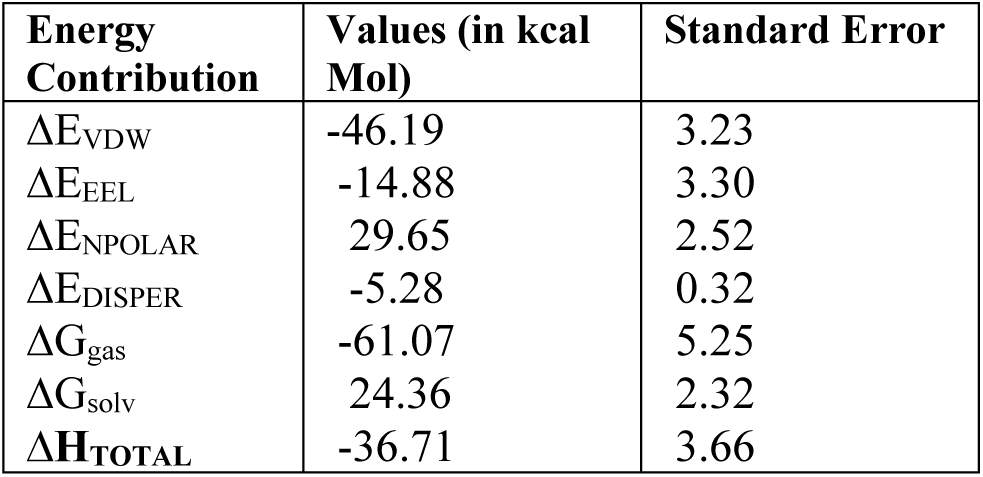
Energy contribution when GP120 bind with NBD in Asn293Gly Mutant. All abbreviations have same meaning as defined in Table 1.

**Table 3.**
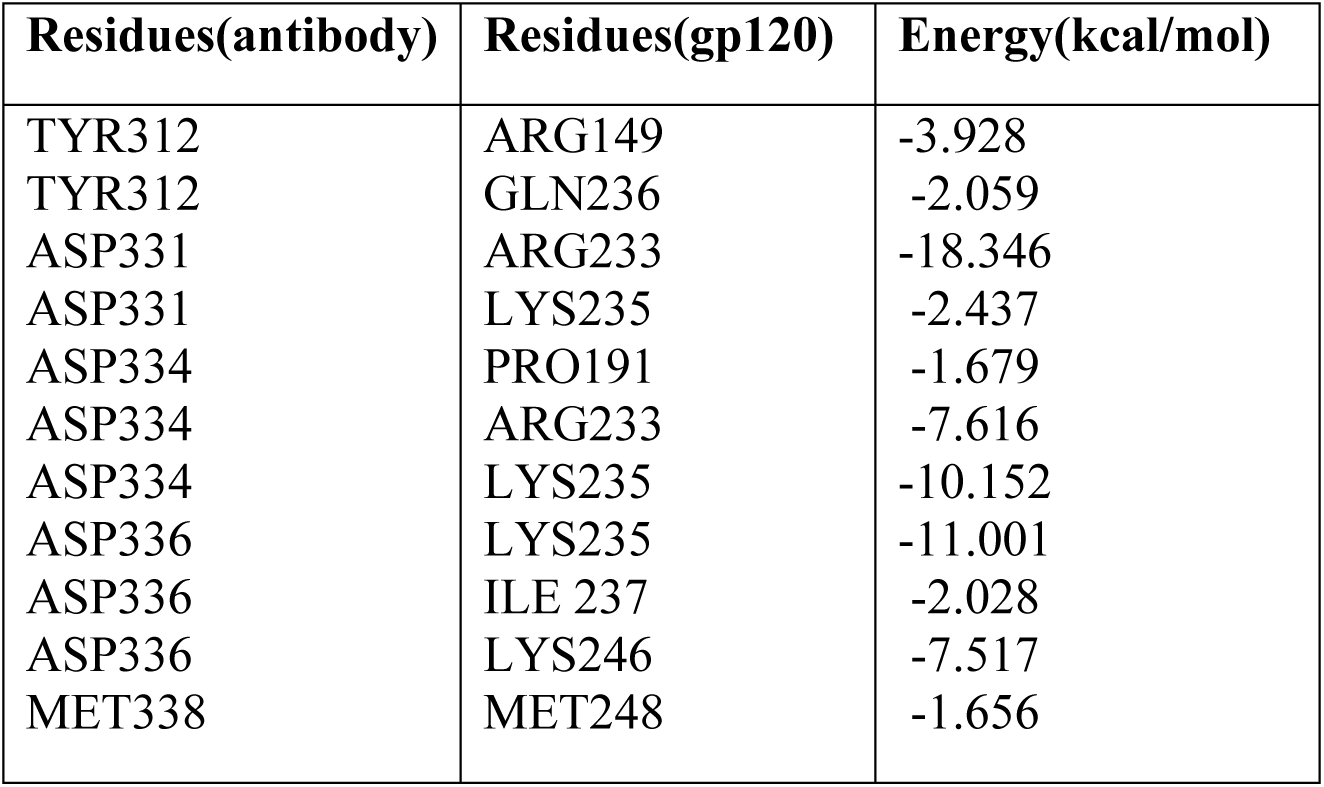
The interaction between antibody and gp120 residues.

| Residues(antibody) | Residues(gp120) | Energy(kcal/mol) |
| --- | --- | --- |
| TYR312 | ARG149 | -3.928 |
| TYR312 | GLN236 | -2.059 |
| ASP331 | ARG233 | -18.346 |
| ASP331 | LYS235 | -2.437 |
| ASP334 | PRO191 | -1.679 |
| ASP334 | ARG233 | -7.616 |
| ASP334 | LYS235 | -10.152 |
| ASP336 | LYS235 | -11.001 |
| ASP336 | ILE 237 | -2.028 |
| ASP336 | LYS246 | -7.517 |
| MET338 | MET248 | -1.656 |

**Table 4.** The thermodynamical parameters for NBD-557 when binds with gp120 in presence of antibody. All abbreviations have same meaning as defined in Table 1

| Energy Contribution | Values (in kcal Mol) | Standard Error |
| --- | --- | --- |
| $\Delta E_{VDW}$ | -43.94 | 2.16 |
| $\Delta E_{EEL}$ | -14.12 | 2.19 |
| $\Delta E_{NPOLAR}$ | 29.40 | 1.46 |
| $\Delta E_{DISPER}$ | -4.90 | 0.15 |
| $\Delta G_{gas}$ | -58.07 | 2.75 |
| $\Delta G_{solv}$ | 24.49 | 1.40 |
| $\Delta H_{TOTAL}$ | **-33.57** | 2.38 |

**Figure 6.**
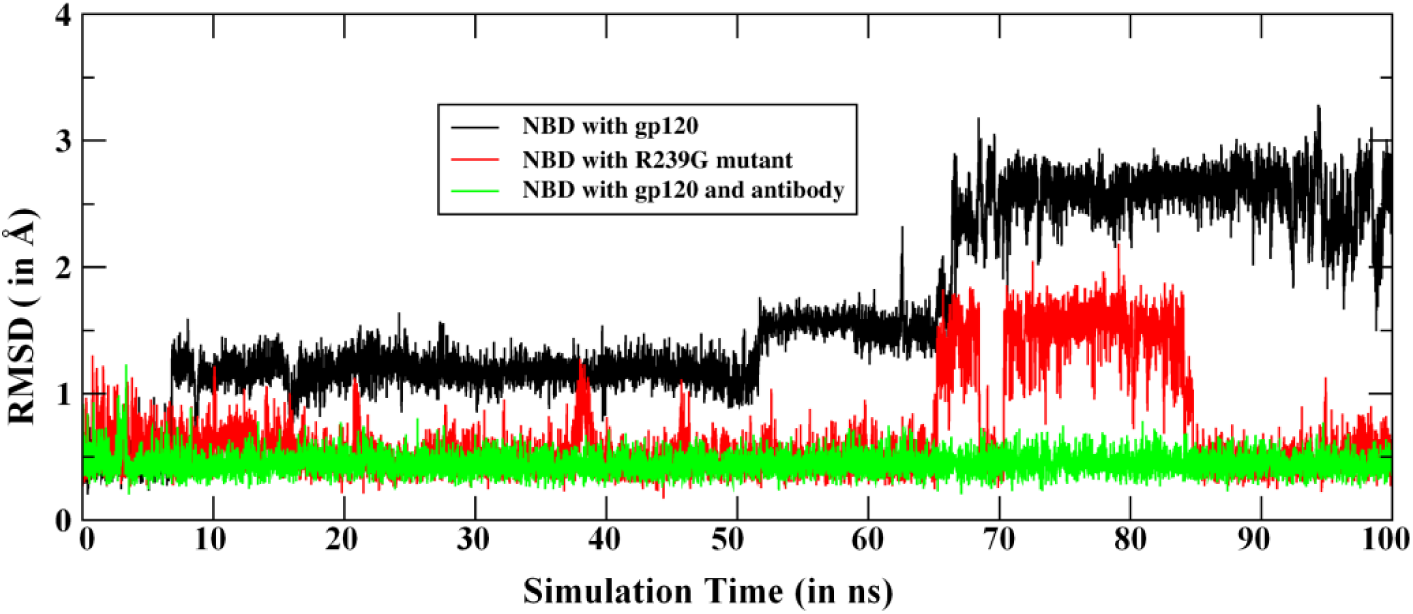
The RMS deviations for NBD-557 in all three MD simulations, i.e. with gp120, in Arg239Gly mutant and with antibody.

**Figure 7.**
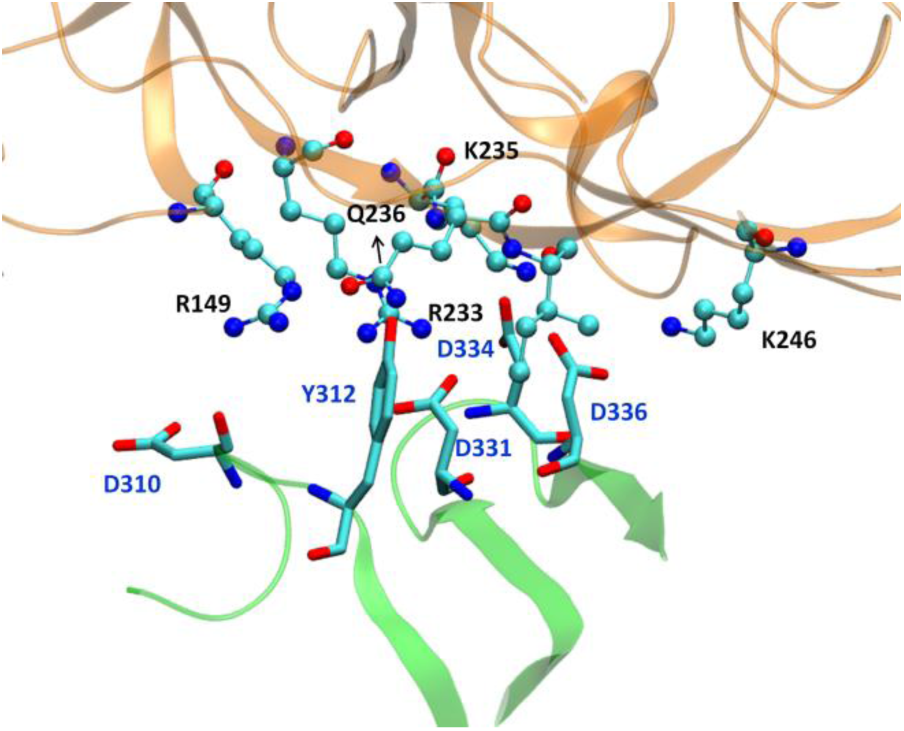
The orientation of interfacial residues of antibody and gp120 protein. The part of gp120 is shown in golden while antibody is shown in light green colour.

**Figure 8.**
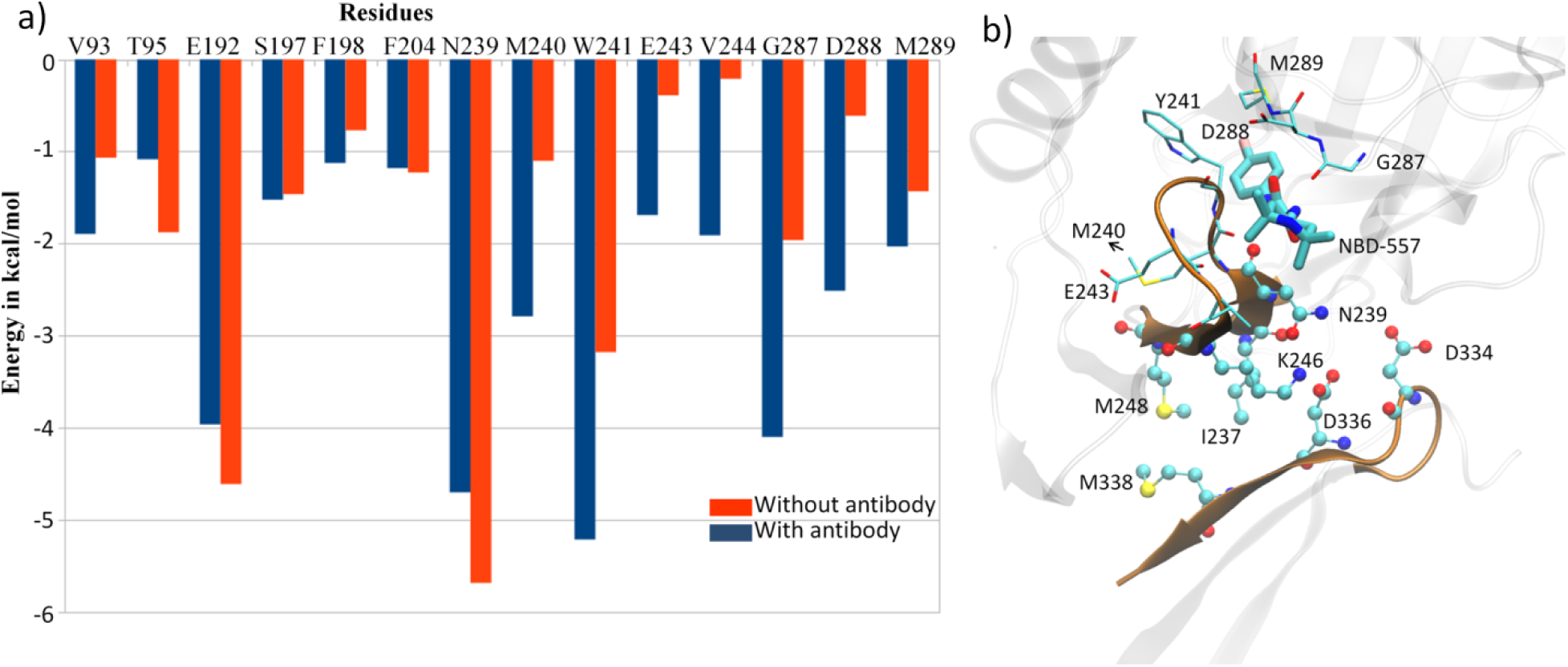
a) A comparative residue interaction map between gp120 and inhibitor NBD-557 in presence and absence of antibody, b) the representation of residues at bridging sheet and antibody.

## 4. Conclusions

NBD-557 is a CD4 mimetic and its competitive receptor which is believed to induce similar conformational changes as CD4 receptor. Our MD simulations clearly show that NBD-557 induces large conformational flexibility, particularly in outer and inner domains. The absence of such flexibility in apo state of gp120 further substantiates the fact that such dynamics is mainly instigated by inhibitor, which was in good agreement with experimental observations. A thorough inspection of the simulated trajectories shows that interaction of NBD-557 with Asn239 is a key factor that choreographs the substrate ligand-induced conformational flexibility in gp120. The role of Asn239 was further supported by the MD simulations of N239G mutant. We have found that the N239G mutation increases the binding affinity but it doesn’t induce the flexibility in gp120 due to lack of flexible side chain interaction with NBD-557. The thermodynamical parameters of NBD-557 in presence of antibody shows enhanced binding enthalpy of inhibitors. We found that antibody possess many acidic residues at its gp120 facing interface which interacts strongly with basic residues of gp120 and thus provide an additional stability to the gp120.NBD-557 complex which in turn, increases the binding affinity of the complex. Our results from the MD simulations of the gp120 with antibody (Fab80) provide a theoretical validation that the antibody stabilizes the gp120 via crucial acid-base pairs at the binding interface between antibody and gp120. The binding free energy calculations quantitatively shows that antibody binding increases change in total binding energy via some conformational rearrangement in bridging sheet β20 which forms the active site for NBD-557 binding.

In summary, present study provide a comprehensive molecular mechanism of NBD-557 binding and how it mimics conformational rearrangement by CD4 co-receptor. Our study provides a theoretical validation of the fact that antibody stabilizes the gp120 and ligand binding. The mechanism of conformational changes instigated by NBD-557 and antibody can be useful for future studies for targeting HIV-1 entry.

## 5. Acknowledgement

RPO, VP and RKT acknowledge Department of Science and Technology (DST), New Delhi for the computational facilities in the form of FIST scheme. KDD acknowledges the Sven Lilly Lawski Fellowship Stockholm, Sweden for the financial support.

## Supporting Information

The SI contains the details of the clustering, details of the residue wise interactions and RMS deviation plots.

## TOC

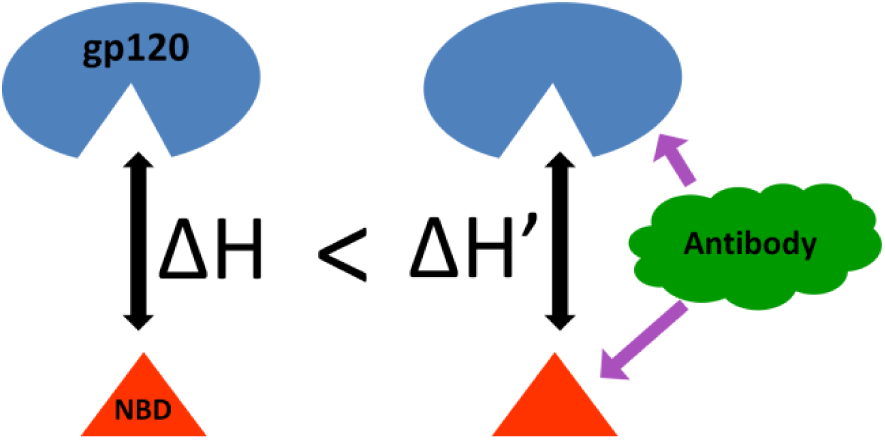

