## Supplementary for "Does Antibody Stabilize the Ligand Binding in GP120 of HIV-1 Envelope Protein? Evidence from MD Simulation"

**Table of Content:**

**Figure S1.** The root mean square deviations of protein backbone, in presence and absence of the inhibitor, NBD-557………………………………… S2

**Figure S2.** The distance between the Glu192 and Asn239 when there is no substrate. We note that H-bond interaction is very week when there is no substrate…… S2

**Table S1.** The thermodynamic parameters for the N239G Mutant……. S2

**Table S2.** Details of residuewise energy decompositions for interaction of NBD-557…. S3

**Table S3**. Details of residuewise energy decompositions for interaction of antibody and gp120…S3

**Table S4-S6.** Details of clustering for gp120 with NBD-557, N239 mutant, and gp120 with NBD in presence of antibody… S4

**Figure S1**. The root mean square deviations of protein backbone, in presence and absence of the inhibitor, NBD-557.

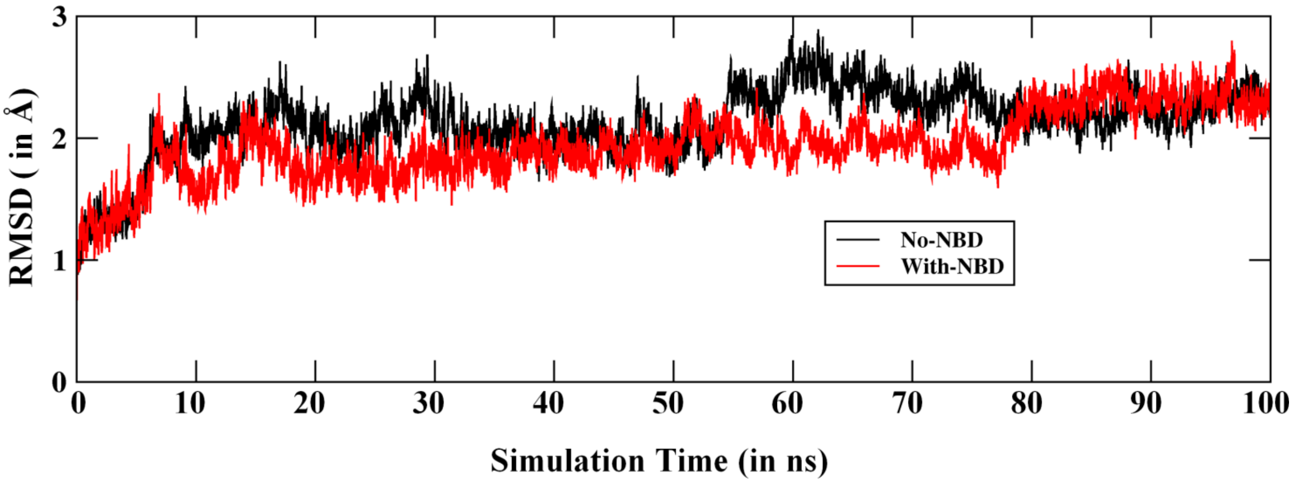


**Figure S2.** The distance between the Glu192 and Asn239 when there is no substrate. We note that H-bond interaction is very week when there is no substrate.

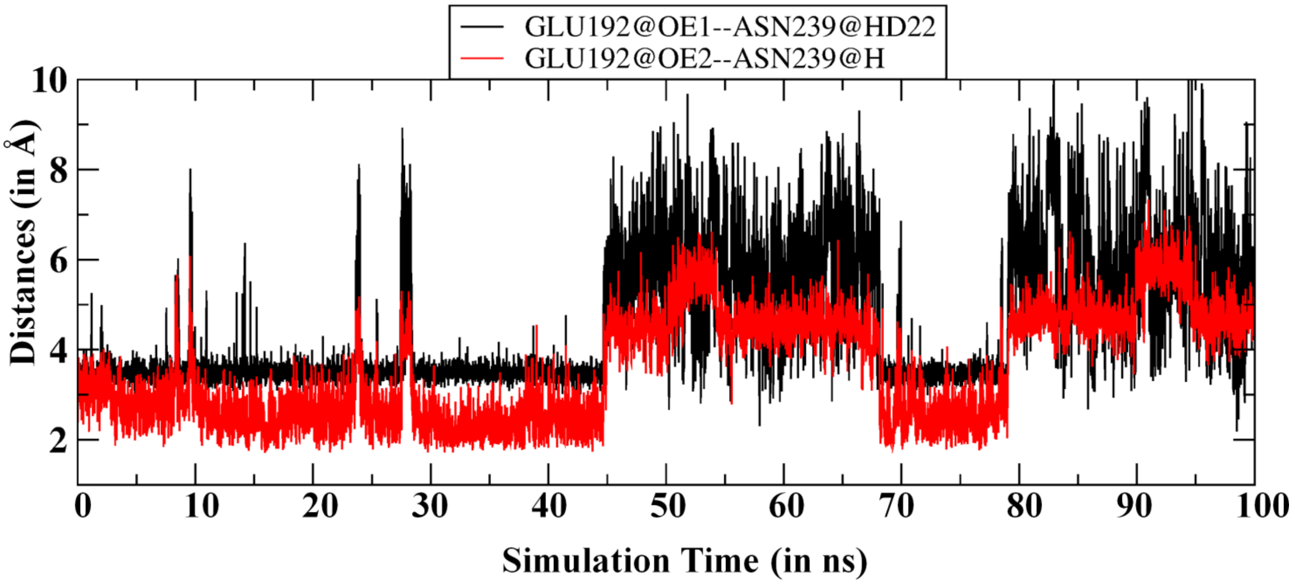


**Table S1.** The thermodynamic parameters for the N239G Mutant.

| **Energy Contribution** | **Values (in kcal Mol)** | **Standard Error** |
|---|---|---|
| $\Delta E_{VDW}$ | -46.19 | 3.23 |
| $\Delta E_{EEL}$ | -14.88 | 3.30 |
| $\Delta E_{NPOLAR}$ | 29.65 | 2.52 |
| $\Delta E_{DISPER}$ | -5.28 | 0.32 |
| $\Delta G_{gas}$ | -61.07 | 5.25 |
| $\Delta G_{solv}$ | 24.36 | 2.32 |
| **$\Delta H_{TOTAL}$** | **-36.71** | 3.66 |

**Table S2.** Details of residuewise energy decompositions for interaction of NBD-557. All energy values are in kcal/mol. Data are shown as value ± standard error.

Total Energy Decomposition:

| Resid 1 | Resid 2 | van der Waals | Electrostatic | Polar Solvation | Non-Polar Solv. | TOTAL |
|---|---|---|---|---|---|---|
| NBD 557 | TRP 23 | -0.800 ± 0.238 | -0.130 ± 0.055 | 0.169 ± 0.054 | -0.492 ± 0.140 | -1.253 ± 0.382 |
| NBD 557 | VAL 93 | -0.658 ± 0.199 | 0.198 ± 0.217 | -0.318 ± 0.103 | -0.291 ± 0.092 | -1.069 ± 0.268 |
| NBD 557 | SER 94 | -0.565 ± 0.136 | -0.360 ± 0.100 | 0.134 ± 0.063 | -0.207 ± 0.063 | -0.998 ± 0.240 |
| NBD 557 | THR 95 | -1.146 ± 0.240 | -0.164 ± 0.188 | 0.249 ± 0.145 | -0.817 ± 0.145 | -1.879 ± 0.379 |
| NBD 557 | ASP 190 | -0.636 ± 0.173 | -0.501 ± 0.328 | 0.242 ± 0.292 | -0.499 ± 0.152 | -1.394 ± 0.357 |
| NBD 557 | PRO 191 | -0.139 ± 0.021 | 0.150 ± 0.039 | -0.115 ± 0.049 | -0.000 ± 0.001 | -0.105 ± 0.034 |
| NBD 557 | GLU 192 | -2.357 ± 0.322 | -1.218 ± 0.550 | 0.432 ± 0.484 | -1.472 ± 0.160 | -4.614 ± 0.610 |
| NBD 557 | ILE 193 | -1.195 ± 0.254 | 0.073 ± 0.062 | -0.054 ± 0.048 | -1.127 ± 0.199 | -2.303 ± 0.398 |
| NBD 557 | HID 196 | -0.072 ± 0.013 | -0.235 ± 0.055 | 0.138 ± 0.051 | 0.000 ± 0.000 | -0.168 ± 0.034 |
| NBD 557 | SER 197 | -0.937 ± 0.209 | 0.054 ± 0.216 | -0.157 ± 0.145 | -0.426 ± 0.087 | -1.465 ± 0.338 |
| NBD 557 | PHE 198 | -0.518 ± 0.131 | -0.229 ± 0.102 | 0.099 ± 0.040 | -0.123 ± 0.039 | -0.771 ± 0.189 |
| NBD 557 | PHE 204 | -0.752 ± 0.228 | -0.283 ± 0.079 | 0.286 ± 0.043 | -0.480 ± 0.076 | -1.229 ± 0.257 |
| NBD 557 | TYR 206 | -0.587 ± 0.142 | -0.242 ± 0.200 | 0.036 ± 0.162 | -0.335 ± 0.081 | -1.128 ± 0.245 |
| NBD 557 | ILE 238 | -0.781 ± 0.153 | 0.360 ± 0.103 | -0.036 ± 0.049 | -0.507 ± 0.108 | -0.964 ± 0.202 |
| NBD 557 | ASN 239 | -2.232 ± 0.399 | -2.285 ± 0.723 | 0.397 ± 0.177 | -1.568 ± 0.148 | -5.688 ± 0.805 |
| NBD 557 | MET 240 | -0.920 ± 0.197 | 0.711 ± 0.489 | -0.476 ± 0.202 | -0.416 ± 0.110 | -1.102 ± 0.498 |
| NBD 557 | TRP 241 | -2.046 ± 0.383 | 0.008 ± 0.218 | 0.117 ± 0.098 | -1.259 ± 0.232 | -3.181 ± 0.514 |
| NBD 557 | GLY 286 | -0.416 ± 0.334 | 0.073 ± 0.298 | 0.022 ± 0.132 | -0.274 ± 0.285 | -0.596 ± 0.611 |
| NBD 557 | GLY 287 | -0.941 ± 0.282 | -0.235 ± 0.282 | 0.123 ± 0.175 | -0.910 ± 0.241 | -1.963 ± 0.515 |
| NBD 557 | MET 289 | -0.779 ± 0.197 | -0.012 ± 0.215 | -0.060 ± 0.116 | -0.584 ± 0.111 | -1.434 ± 0.248 |

**Table S3**. Details of residuewise energy decompositions for interaction of antibody and gp120. All energy values are in kcal/mol. Data are shown as value ± standard error.

Total Energy Decomposition:

| Resid 1 | Resid 2 | van der Waals | Electrostatic | Polar Solvation | Non-Polar Solv. | TOTAL |
|---|---|---|---|---|---|---|
| ASP 310 | GLN 150 | -0.208 ± 0.194 | -2.305 ± 2.614 | 2.115 ± 2.231 | -0.182 ± 0.227 | -0.579 ± 0.919 |
| TYR 312 | ARG 149 | -1.448 ± 0.332 | -1.108 ± 0.808 | -0.166 ± 0.578 | -1.207 ± 0.254 | -3.928 ± 0.832 |
| TYR 312 | LYS 235 | -0.222 ± 0.068 | 1.205 ± 0.484 | -1.765 ± 0.404 | -0.127 ± 0.048 | -0.909 ± 0.282 |
| TYR 312 | GLN 236 | -1.012 ± 0.244 | 0.150 ± 0.485 | -0.446 ± 0.316 | -0.751 ± 0.144 | -2.059 ± 0.450 |
| LEU 329 | ILE 237 | -0.284 ± 0.104 | -0.033 ± 0.013 | 0.066 ± 0.016 | -0.352 ± 0.098 | -0.604 ± 0.164 |
| ASP 331 | ARG 233 | 1.160 ± 0.969 | -51.864 ± 2.007 | 32.894 ± 1.245 | -0.536 ± 0.035 | -18.346 ± 1.813 |
| ASP 331 | LYS 235 | -0.204 ± 0.050 | -29.633 ± 1.672 | 27.487 ± 1.239 | -0.087 ± 0.033 | -2.437 ± 0.651 |
| GLU 333 | ARG 233 | -0.984 ± 0.314 | -22.713 ± 2.701 | 23.804 ± 2.301 | -1.013 ± 0.135 | -0.905 ± 0.811 |
| ASP 334 | PRO 191 | -0.731 ± 0.194 | -2.548 ± 0.427 | 2.206 ± 0.355 | -0.605 ± 0.134 | -1.679 ± 0.374 |
| ASP 334 | ARG 233 | -0.011 ± 0.621 | -36.558 ± 2.139 | 29.505 ± 0.810 | -0.553 ± 0.062 | -7.616 ± 1.396 |
| ASP 334 | LYS 235 | 0.256 ± 0.708 | -43.911 ± 2.874 | 34.074 ± 1.058 | -0.571 ± 0.068 | -10.152 ± 1.845 |
| ASP 336 | ARG 233 | -0.087 ± 0.018 | -22.654 ± 0.875 | 21.651 ± 0.640 | -0.019 ± 0.011 | -1.109 ± 0.337 |
| ASP 336 | LYS 235 | 0.507 ± 0.620 | -47.018 ± 1.939 | 35.887 ± 0.752 | -0.376 ± 0.037 | -11.001 ± 1.200 |
| ASP 336 | ILE 237 | -1.215 ± 0.244 | 2.300 ± 0.830 | -2.176 ± 0.582 | -0.937 ± 0.143 | -2.028 ± 0.423 |
| ASP 336 | LYS 246 | -0.050 ± 0.631 | -42.962 ± 3.866 | 36.184 ± 1.825 | -0.690 ± 0.136 | -7.517 ± 2.089 |
| MET 338 | VAL 31 | -0.531 ± 0.203 | -0.033 ± 0.143 | 0.042 ± 0.110 | -0.538 ± 0.175 | -1.060 ± 0.359 |
| MET 338 | MET 248 | -0.797 ± 0.256 | -0.192 ± 0.220 | 0.065 ± 0.093 | -0.732 ± 0.191 | -1.656 ± 0.441 |

**Table S4.** Clustering of gp120 with NBD without antibody

| #Cluster | Frames | Frac | AvgDist | Stdev | Centroid | AvgCDist |
|---|---|---|---|---|---|---|
| 0 | 2565 | 0.218 | 1.636 | 0.208 | 3429 | 2.281 |
| 1 | 1912 | 0.127 | 1.591 | 0.202 | 6565 | 2.560 |
| 2 | 1520 | 0.121 | 1.688 | 0.237 | 1663 | 2.357 |
| 3 | 824 | 0.115 | 1.591 | 0.215 | 6002 | 2.515 |
| 4 | 729 | 0.102 | 1.599 | 0.198 | 2063 | 2.378 |
| 5 | 566 | 0.079 | 1.516 | 0.219 | 1146 | 2.349 |
| 6 | 532 | 0.074 | 1.559 | 0.199 | 261 | 2.618 |
| 7 | 502 | 0.070 | 1.572 | 0.215 | 5254 | 2.301 |
| 8 | 379 | 0.053 | 1.524 | 0.204 | 4824 | 2.257 |
| 9 | 295 | 0.041 | 1.520 | 0.246 | 684 | 2.419 |

**Table S5.** Clustering of gp120 with NBD in N239G mutant.

| #Cluster | Frames | Frac | AvgDist | Stdev | Centroid | AvgCDist |
|---|---|---|---|---|---|---|
| 0 | 1803 | 0.180 | 1.578 | 0.164 | 3230 | 2.037 |
| 1 | 1569 | 0.157 | 1.528 | 0.174 | 7695 | 2.173 |
| 2 | 1512 | 0.151 | 1.551 | 0.177 | 6321 | 2.142 |
| 3 | 1368 | 0.137 | 1.533 | 0.172 | 9362 | 2.120 |
| 4 | 1040 | 0.104 | 1.497 | 0.151 | 5347 | 2.089 |
| 5 | 731 | 0.073 | 1.535 | 0.189 | 2134 | 2.050 |
| 6 | 692 | 0.069 | 1.600 | 0.210 | 780 | 2.225 |
| 7 | 511 | 0.051 | 1.511 | 0.194 | 1689 | 2.107 |
| 8 | 454 | 0.045 | 1.531 | 0.199 | 328 | 2.429 |
| 9 | 320 | 0.032 | 1.519 | 0.195 | 1365 | 2.158 |

**Table S6**. Clustering of gp120 with NBD in presence of antibody.

| #Cluster | Frames | Frac | AvgDist | Stdev | Centroid | AvgCDist |
|---|---|---|---|---|---|---|
| 0 | 4934 | 0.493 | 2.508 | 0.523 | 4357 | 10.009 |
| 1 | 1370 | 0.137 | 2.621 | 0.632 | 7213 | 5.879 |
| 2 | 963 | 0.096 | 2.731 | 0.843 | 5298 | 7.652 |
| 3 | 900 | 0.090 | 2.597 | 0.689 | 9238 | 5.369 |
| 4 | 473 | 0.047 | 2.130 | 0.518 | 8553 | 6.321 |
| 5 | 367 | 0.037 | 2.230 | 0.642 | 9822 | 6.619 |
| 6 | 310 | 0.031 | 2.300 | 0.668 | 9471 | 6.176 |
| 7 | 306 | 0.031 | 1.918 | 0.408 | 6263 | 5.332 |
| 8 | 283 | 0.028 | 1.909 | 0.478 | 6013 | 5.813 |
| 9 | 94 | 0.009 | 1.677 | 0.416 | 6830 | 6.540 |
